# Fungi with history: Unveiling the mycobiota of historic documents of Costa Rica

**DOI:** 10.1101/2022.06.12.495835

**Authors:** Efraín Escudero-Leyva, Sofía Vieto, Roberto Avendaño, Diego Rojas-Gätjens, Paola Agüero, Carlos Pacheco, Mavis L. Montero, Priscila Chaverri, Max Chavarría

## Abstract

Through nondestructive techniques, we studied the physicochemical characteristics and mycobiota of five key historic documents from Costa Rica, including the Independence Act of Costa Rica from 1821. We determined that for documents dated between 1500 and 1900 (i.e., the Cloudy Days Act, the Independence Act, and two documents from the Guatemalan Series from 1539 and 1549), the paper composition was cotton, whereas the 1991 replicate of the Political Constitution from 1949 was made of wood cellulose with an increased lignin content. We also determined that the ink employed in 1821 documents is ferrogallic, i.e., formed by iron sulfate salts in combination with gallic and tannic acids. In total, 22 fungal isolates were obtained: 15 from the wood-cellulose-based Political Constitution and seven from the other three cotton-based documents. These results suggest that cotton-based paper is the most resistant to microbial colonization. Molecular identifications using three DNA markers (i.e., ITS nrDNA, beta-tubulin, and translation elongation factor 1-alpha) classified the isolates in eight orders and ten genera. The most frequent genera were *Cladosporium, Penicillium*, and *Purpureocillium*. Of the isolates, 95% presented cellulolytic activity correlated to their ability to cause deterioration of the paper. This work increases the knowledge of the fungal diversity that inhabits historic documents and its relationship with paper composition and provides valuable information to develop strategies to conserve and restore these invaluable documents.

## Introduction

Because biodeterioration can lead to the damage of historic documents, artwork, monuments, or buildings, its study is fundamental for the conservation of cultural heritage (Sterflinger and Pinar 2012; Palla and Barresi 2017; Vieto *et al*. 2021; Ranalli and Zanardini 2021; Ranalli *et al*. 2005). The prevention of biodeterioration and development of adequate conservation and restoration strategies cannot be an unscripted process; it is necessary to undertake diagnoses of these valuable pieces of our history and art, which include chemical characterization and the study of microbial diversity together with the physiological characteristics of biodeteriogens (Palla and Barresi 2017; Negi and Sarethy 2019).

Valuable cultural and historic objects, such as relevant paintings, ancient sculptures, and historic documents, can be seen as substrates on which microorganisms can thrive and cause damage. Specifically, paper-based documents contain biodegradable organic constituents that fungi can use as a substrate (Jia *et al*. 2020; Pyzik *et al*. 2021). The term “paper” is a general concept that encompasses all thinly laminated material that is produced with vegetable fiber pulp or other materials ground and mixed with water, dried, and hardened. Historically, vegetable fibers have been extracted from natural sources, such as straw, silk, hemp, flax, cotton, and the bark of different trees, among others. The content of cellulose and other components of paper can vary depending on its origin, generating papers that are more or less resistant to biodegradation as a consequence. For example, it is well known that cellulose fibers have high purity in cotton and linen papers, which results in papers with greater durability and resistance to biodeterioration (Negulescu *et al*. 1998; Daria *et al*. 2020).

Damage produced by fungi that is normally present on paper —including staining, material weakening, and partial or complete destruction of documents— can occur in the long term (Sequeira *et al*. 2019). Besides the alterations caused to the documents, the health of curators and people involved in archives or museums can also be threatened if the spore production is elevated or mycotoxins are produced (Sterflinger and Pinzari 2012). The study of fungi responsible for the biodegradation of paper began in 1818 with the pioneering work of Christian Gottfried Ehrenberg (Sterflinger and Pinzari 2012). To date, diverse fungi have been identified in old paperwork, such as *Aspergillus, Chaetomium, Cladosporium, Penicillium*, and *Trichoderma;* occasionally, new species can even be found (Coronado-Ruiz *et al*. 2018; Sequeira *et al*. 2019). Mesquita *et al*. (2009) isolated, identified, and characterized the microbiota from historic documents dated between 1860–1939 in the archive of the University of Coimbra and found fourteen fungal genera, of which *Aspergillus*, *Cladosporium*, and *Penicillium* were the most common. Our research group recently isolated nineteen fungi from a nineteenth-century French collection of drawings and lithographs in the custody of Universidad de Costa Rica (Coronado-Ruíz *et al*. 2018). The fungi were molecularly identified as *Arthrinium, Aspergillus*, *Chaetomium*, *Cladosporium*, *Colletotrichum*, *Penicillium*, and *Trichoderma*; a great majority of them showed cellulolytic activity. Many fungal species found in historic paper-based documents contain enzymatic activity related to biodeterioration, which allows fungi to use these surfaces as a source of carbon (Coronado-Ruíz *et al*. 2018; Pinheiro *et al*. 2019; Vieto *et al*. 2022). The enzymatic machinery to take advantage of paper as a source of carbon has been reported in fungi isolated from historic documents and includes the presence of exoenzymes with cellulase activity (Puškárová *et al*. 2019; El Begardi *et al*. 2014), lignocellulolytic (Mazzoli *et al*. 2018) glucanase, and laccase (Sterflinger and Pinzari 2012).

The National Archive of Costa Rica (NACR) —called Archivos Nacionales (National Archives) before 1948—is where the most treasured documents in Costa Rica are preserved; these include the Cloudy Days Act (Acta de los Nublados, September 28, 1821), in which authorities of the Municipality of León, in the Captaincy General of Guatemala, expressed their position on Central American Independence; the Political Constitution with all historic changes, including the abolition of the Costa Rican army (Jaén García 2019; Chacón León 2021); and perhaps the most important historic document in the country: the Costa Rican Independence Act (Acta de Independencia, October 29, 1821). These invaluable documents are in addition to more than 20,000 linear meters of other paper-based documents that contain the history of Costa Rica and that are in the custody of NACR. Due to the tropical peculiarities of the country—such as high humidity, heat, long rainy seasons, and sometimes inadequate storage conditions—documents and artworks are constantly threatened and come under continuous biodeterioration, making the conservation of the country’s cultural heritage a challenge (Silva, 2011).

The aim of this work was to characterize the paper composition and evaluate the presence and cellulolytic activity of culturable fungi in historic documents from the National Archive of Costa Rica, including invaluable documents such as the Independence Act of Costa Rica. This information enables restorers to establish guidelines for the preservation and restoration of paper-based historic documents.

## Materials and Methods

### Sampling of documents

Permits to sample the historic documents were obtained from the Institutional Commission of Biodiversity of the University of Costa Rica (resolution N° 186) and authorities of the NACR. Historic documents stored at NACR were sampled between March and September 2019. The documents were: (i) Political Constitution redacted in 1949 (1991 replicate), (ii) Cloudy Days Act from 1821 (Acta de los Nublados), (iii) Independence Act of Costa Rica from 1821, and (iv) two documents from the Guatemalan Series from 1539 and 1549 (Fig. 1). For fungal isolation, careful rubbing with sterile cotton swabs over the surface of the documents was performed, especially seeking signs of biodeterioration such as dark spots. The swabs were then saved inside Falcon tubes for transport to the laboratory.

**Figure 1.**
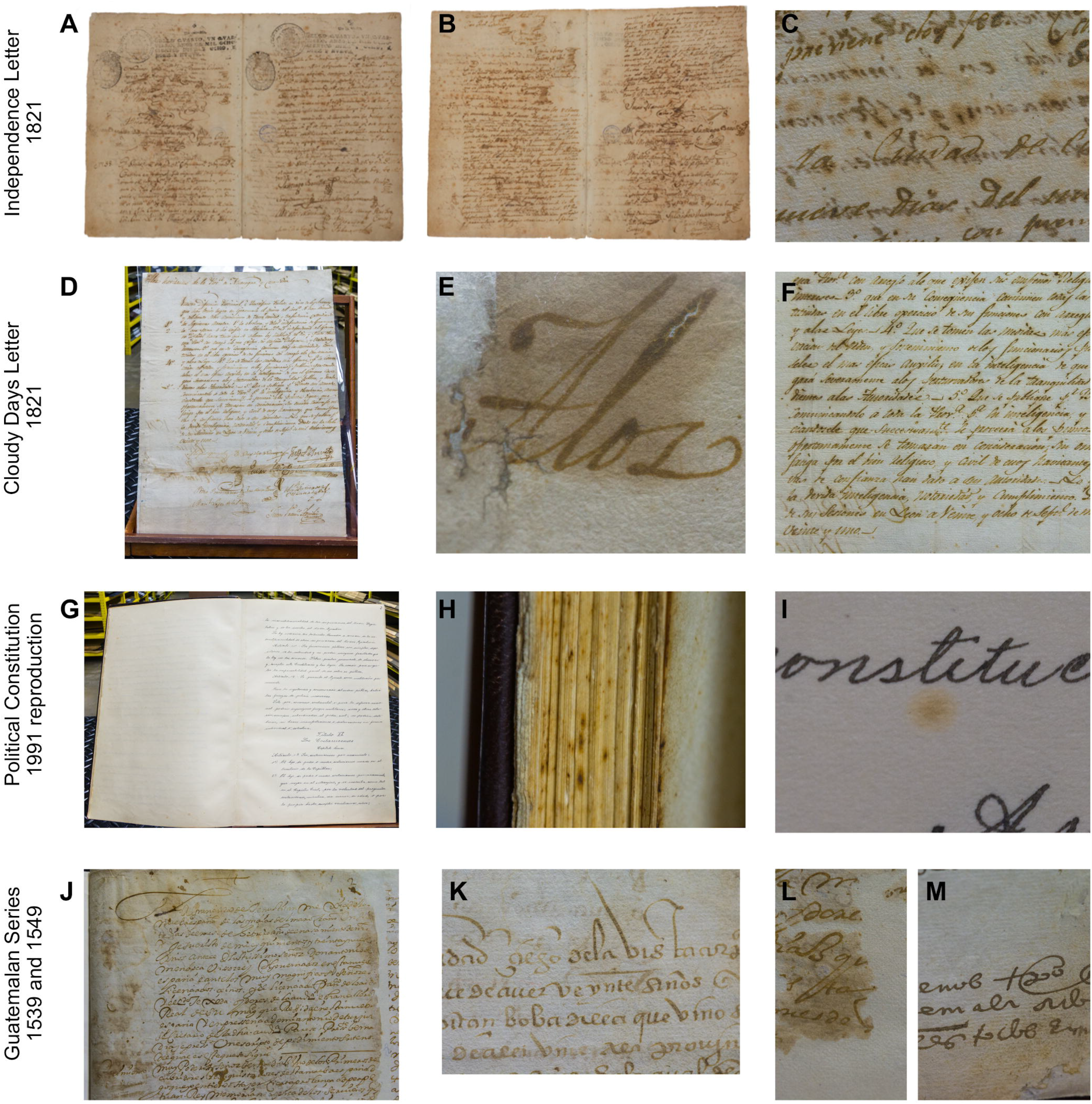
Historic documents from Costa Rica analyzed for chemical and substrate characterization. **A,B**. Independence Act. **C**. Signs of deterioration on the Independence Act, including yellow spots around the letters. **D**. Cloudy Days Act. **E**. Humidity mark on Cloudy Days Act. **F**. Oxidation signs around the ink from Cloudy Days Act. **G**. 1949 Political Constitution (1991 reproduction). **H**. Signs of microbiological contamination on 1949 Political Constitution (1991 reproduction). **I**. Yellow spots on 1949 Political Constitution (1991 reproduction). **J**. Fragment of the 1539 Guatemalan Series. **K**. Fragment of the 1549 Guatemalan Series. **L,M**. Signs of leakage and humidity on 1539-1549 Guatemalan Series.

### Material characterization by attenuated-total-reflection Fourier-transform infrared spectra (ATR-FTIR)

ATR-FTIR was used to determine functional groups and distinguish cellulosic materials. These ATR-FTIR spectra were recorded using a portable spectrophotometer (Bruker Alpha II, Canada) with platinum ATR mode and monolithic diamond crystal. The spectral resolution was 4 cm^-1^, in wavenumber range 400–4,000 cm^-1^ with 99 scans. To carry out the identification, the “Database of ATR-FT-IR spectra of various materials” (Vahur *et al*. 2016) was used.

### Material characterization by X-ray fluorescence (XRF)

X-ray fluorescence (XRF) was used to determine the elemental composition of the material, especially the presence of metallic ions, such as iron and calcium, among others. This technique is especially important in the characterization of inks. The XRF spectra were recorded with a portable XRF spectrophotometer (Elio, XGLab; Bruker, Italy) measured at the Ka line of manganese (resolution 140 eV), a SDD detector (active area 25 mm^2^, fluorescence angle 63°, incident angle 90°) and a distance of 14 mm from the detector to the sample. The electric current was adjusted to 80 μA, with a voltage of 50 kV and a measuring duration of 300 s. Software XRS-FP2 (CrossRoads Scientific, USA) was used for data analysis, maintaining a noise signal of 0.5.

### Fungal cultivation strategy

Samples were processed in the laboratory 2-4 hours after sampling. Cotton swabs were immersed into sterile phosphate-buffered saline solution (PBS; 400 μL, 1X; Thermo Fisher Scientific, USA) and homogenized using a vortex (40 s). Each sample (100 μL) was then cultured in plates of potato dextrose agar (1% Difco PDA; BD company, France) and carboxymethyl cellulose (1% CMC; Sigma Aldrich, USA, with 0.8 % agar; BD company, France) with kanamycin (50 μg/mL; Sigma-Aldrich, USA) and incubated (25 °C) until growth was observed. Colonies exhibiting varied morphologies were purified and transferred onto PDA plates; photographs were taken after incubation (15 days, 30 °C).

### Molecular identification of the isolated fungi

Genomic DNA was extracted from the isolated fungi using the method described by Lodhi *et al*. (1994) with modifications. First, two agar disks (diameter 8 mm) from each fungus were added to a centrifuge tube (2 mL) and were ground with sterile micro-pestles. Extraction buffer (750 μL, sodium EDTA 20 mM, tris-HCl 100 mM, NaCl 1.4 M, CTAB 2 % [w/v], PVP 2% [w/v] and β-mercaptoethanol 0.2%) was added; the tubes were vortexed and incubated (20 min, 65 °C). For DNA separation, trichloromethane-octanol (750 μL, 24:1) was added to the mixture and centrifuged (25 °C, 14,000 rpm). DNA from the top aqueous phase (600 μL) was precipitated on an addition of 2-propanol (600 μL; Sigma-Aldrich, USA). Then, ethanol (70 %, 500 μL; Sigma-Aldrich, USA) was used to wash the precipitated DNA. Finally, DNA was resuspended in Tris-EDTA buffer (50 μL) with RNAse (1 μL, 10 mg/mL; Thermo Fisher Scientific, USA).

To obtain a preliminary identification of the isolates, the nrDNA internal transcribed spacers (ITS) were amplified with primers ITS4 (TCCTCCGCTTATTGATATGC) and ITS5 (GGAAGTAAAAGTCGTAACAAGG) (White *et al*. 1990). Depending on the results from ITS, secondary markers were used to refine the identifications for some isolates, i.e., portions of the translation elongation factor 1-alpha (TEF1; primers CATCGAGAAGTTCGAGAAGG and TACTTGAAGGAACCCTTACC) (Carbone and Kohn 1999) and beta-tubulin (TUB2; primers AACATGCGTGAGATTGTAAGT and TAGTGACCCTTGGCCCAGTTG) (O’Donnell and Cigelnik 1997) genes. Each reaction (total volume 20 μL) consisted of Master Mix (10 μL, 2X; Thermo Fisher Scientific, USA), bovine serum albumin (BSA, 0.5 μL, 20 mg/mL; Sigma Aldrich, USA), dimethyl sulfoxide (DMSO, 1.5 μL; Sigma Aldrich, USA), and primers (0.5 μL each, 10 μM) and DNA (2 μL, 50 ng/μL). PCR reactions were implemented in a thermal cycler (9902 Veriti, Applied Biosystem, Norwalk, USA), according to conditions described by Schoch *et al*. (2012) for ITS, Carbone and Kohn (1999) for TEF1 and O’Donnell and Cigelnik (1997) for TUB2. Sanger sequencing of PCR products was performed with Psomagen (USA); the raw sequences were edited and assembled in Bioedit v.7.2. Isolate identification was performed by comparing the consensus sequences against the GenBank database using the BLAST search tool. Then, a cladogram was constructed using the ITS sequences. For this, the two closest matches, with type material prioritized, were retrieved/downloaded from the BLAST analysis and aligned using MUSCLE (Edgar 2004). The resulting alignment in Phylip format was submitted to Bayesian Inference analysis with Exabayes (Aberer *et al*. 2014). MCMC was run in parallel and 15 million generations were done with 25% burn-in. All analyses were run in the Kabré supercomputer (CNCA-CONARE, Costa Rica). A consensus tree was visualized and edited with FigTree v.1.4.3 (Rambaut 2010). Newly generated sequences were deposited in GenBank under accession numbers ON479855-ON479876 (ITS), ON720280-ON720285 (TEF1), and ON734081–ON734096 (TUB2).

### Screening of cellulolytic activity

The screening of cellulase-producing fungi was undertaken on carboxymethyl cellulose plates (CMC, 1 %; Sigma Aldrich, USA) as the sole carbon source, supplemented with agar (0.8 %, Sigma Aldrich, USA; Johnsen and Krause 2014; Gohel *et al*. 2014). For this purpose, agar disks (diameter ~0.8 mm) of each fungus were placed in the center of CMC plates and incubated (7 days, 30 °C). After incubation, each plate was flooded with Gram’s iodine stain (10 mL; Sigma-Aldrich, USA; Kasana *et al*. 2008; Gohel *et al*. 2014) and washed with water for 10 min. Because Gram’s iodine dye is held only by integral cellulose polymers, cellulase activity is revealed by the clear zones appearing as pale halos (Florencio *et al*. 2012). Photographs were taken before and after staining the plates; software ImageJ v.1.52k (Bourne 2010) was used to measure fungal growth (as the diameter of the colony) and the halo diameter for a subsequent calculation of the enzymatic index (EI), a semiquantitative estimate of enzyme activity according to the following formula (Florencio *et al*. 2012):

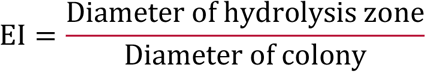

The experiments were performed in triplicate; *Pleurotus ostreatus* served as a positive control (Garzlllo *et al*. 1994; Valášková and Baldrian 2006).

## Results

### Material characterization by infrared spectra (ATR-FTIR) and X-ray fluorescence (XRF)

Macroscopically all documents analyzed showed detailed damage as observed in Fig. 1, in which signs of humidity and possible leakage are visible in several areas. Particularly, in the Independence Act surface, orangish spots were present. When observed under UV light, those areas are fluorescent and appear as dark spots in UV reflectance photographs (Supplementary Fig. S1).

The chemical composition of historic documents (both inks and paper) was studied through nondestructive and portable techniques (See Materials and Methods). The composition of each document is shown in Table 1. The composition of the organic substrate was determined by comparing the IR spectra with databases; the elements present in the ink and additives were resolved with X-ray fluorescence. The results showed that the paper in the documents from 1500–1900 was handmade mainly from cotton with watermark presence (excepting the second page from the Cloudy Days Act, which was cellulose-based). In the Political Constitution (1991 replicate), modern paper was used, characterized by shorter fibers and greater lignin content; this indicates that this paper was made from wood cellulose. The results showed that the ink employed in the documents from 1821 was ferrogallic, formed by iron sulfate salts in combination with gallic and tannic acids (Table 1 and Supplementary Fig. S1).

**Table 1.** Chemical, substrate, and ink characterization of the historic documents from Costa Rica.

| Document | Sections analyzed for chemical characterization | Sections analyzed for fungal isolation | Paper composition | Ink chemical elements detected |
| --- | --- | --- | --- | --- |
| Independence Act, 1821 | Pages 1-3 | Pages 1-3 | Cotton | Fe, Ca, Zn, K |
| Cloudy Days Act, 1821 | Page 1<br>Page 2 | Full document | Cotton<br>Cellulose acetate | Fe, Ca, Zn, K, S, Cl, Pb<br>Fe, Ca, Zn |
| Political Constitution, 1949 (1991 reproduction) | Full document | Cover and Slavery Abolition page | Wood cellulose | Not determined |
| Guatemalan Series 1539 | Full document | Pages 1,8,3,17 | Cotton | Not determined |
| Guatemalan Series 1549 | Full document | Pages 2,5,9,10 | Cotton | Not determined |

### Isolation and identification of fungi

In total, 22 fungal isolates (Supplementary Fig. S2) were recovered from the Costa Rican historic documents, being the Political Constitution (1991 replicate) that with the most isolates (15 in total). The taxonomic identification provided by BLAST and ITS phylogenetic analyses are shown in Table 2 and Fig. 2, respectively. The fungi recovered belong to 14 genera and eight orders. Two taxa were obtained from the Independence Act, two from the Cloudy Days Act, two from the document from 1549, and only one fungal isolate from the oldest document (Guatemalan Series 1539). The phylogenetic placement of the isolates corresponded to eight orders, as shown in the ITS cladogram (Fig. 2). The phylogenetic analysis supports the identifications performed with the BLAST tool, at least at the genus level. TEF1 and TUB2 sequences refined the identification for some of the isolates (Table 2).

**Figure 2.**
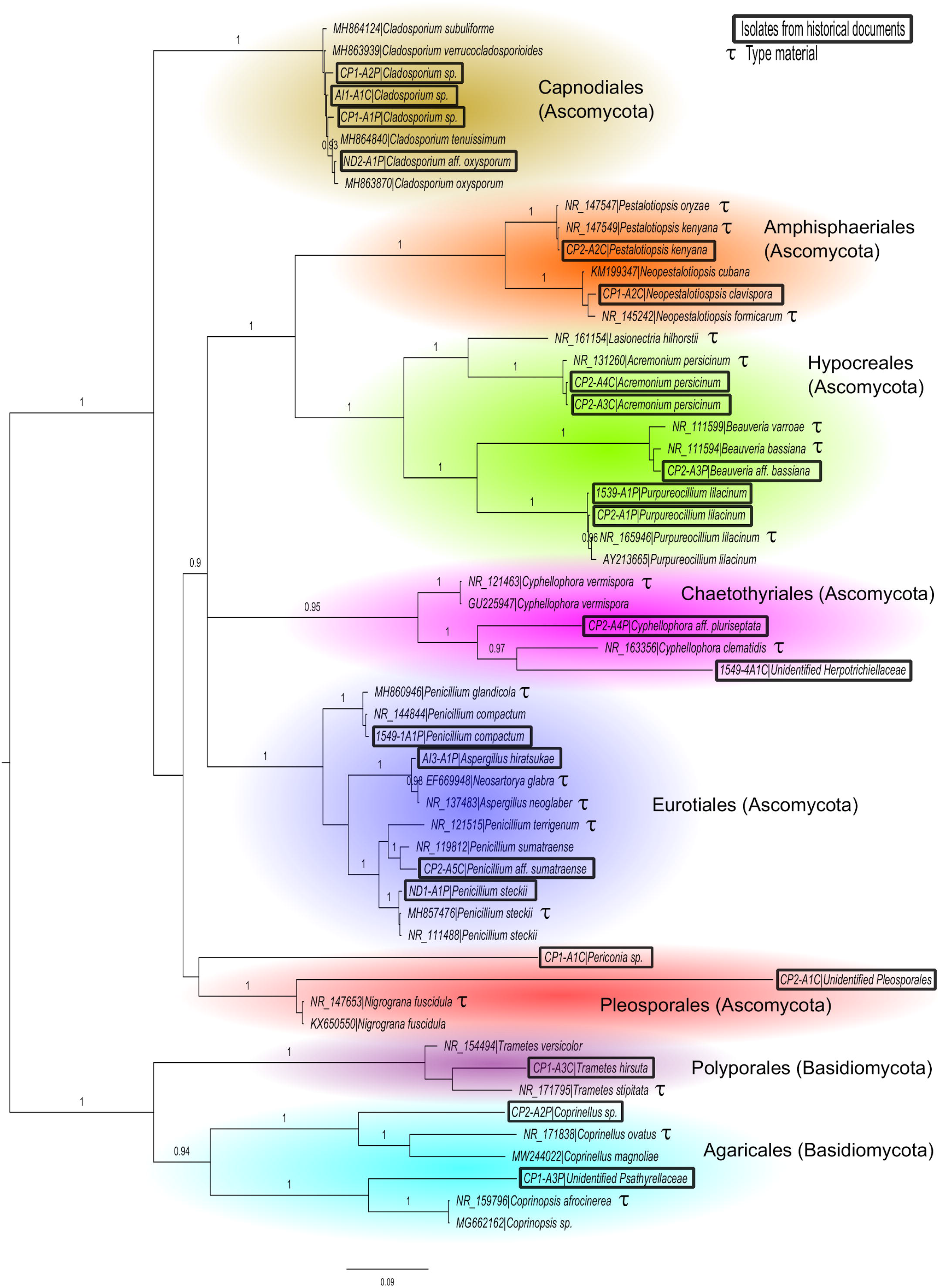
Bayesian Inference consensus cladogram based on nrDNA ITS sequences. Posterior probabilities are indicated at branches. LnL = −13564.21

**Table 2.** Identification (ID); BLAST results using ITS, TEF1 and TUB2; and origin of documents. Names in bold indicate the assigned classification.

| Isolate number | Origin | Closest match and accession number / % similarity * |  |  |
| --- | --- | --- | --- | --- |
|  |  | ID with ITS | ID with TEF1 | ID with TUB2 |
| 1539-A1P | Guatemalan series, 1539 | ***Purpureocillium lilacinum***<br>MZ359582 / 99.3 | ***Purpureocillium lilacinum***<br>MH613753 / 100 | ***Purpureocillium lilacinum***<br>JQ965112 / 99.8 |
| 1549-1A1P | Guatemalan series, 1549 | ***Penicillium* sp.** FJ752622 / 99.81 | - | ***Penicillium compactum***<br>KM973202 / 98.6 |
| 1549-4A1C |  | **Unidentified**<br>**Herpotrichiellaceae**<br>KJ612089 / 90.48 | no match | no match |
| AI1-A1C | Independence Act, 1821 | ***Cladosporium* sp.**<br>OK274323 / 100 | - | ***Cladosporium oxysporum***<br>EF101455 / 95.5 |
| AI3-A1P |  | ***Aspergillus hiratsukae***<br>MN347034 / 100 | - | ***Aspergillus hiratsukae***<br>MH644026 / 100 |
| CP1-A1C | Political Constitution,<br>1949 (1991 reproduction) | ***Periconia* sp.** KP128003 / 99.80 | no match | no match |
| CP1-A1P |  | ***Cladosporium* sp.**<br>OK274323 / 92.48 | - | ***Cladosporium* sp.**<br>JQ217373 / 97 |
| CP1-A2C |  | ***Pestalotiopsis microspora***<br>OK254042 / 100 | - | ***Neopestalotiopsis***<br>***clavispora*** OM328818 / 98.2 |
| CP1-A2P |  | ***Cladosporium* sp.**<br>EF504401 / 100 | - | - |
| CP1-A3C |  | ***Trametes hirsuta***<br>GQ280373 / 100 | - | - |
| CP1-A3P |  | **Unidentified**<br>**Psathyrellaceae** JQ922137 / 100 | - | no match |
| CP2-A1C |  | **Unidentified Pleosporales**<br>KP263091 / 92.42 | - | ***Biatrispora* sp.** MF588919 / 86 |
| CP2-A1P |  | *Purpureocillium lilacinum*<br>MZ359582 / 99.88 | - | *Purpureocillium lilacinum*<br>GU968702 / 100% |
| CP2-A2C |  | *Pestalotiopsis trachycarpicola* MZ453106<br>/ 99.82 | *Neopestalotiopsis* sp.<br>KR493607 / 100 | *Pestalotiopsis kenyana*<br>KX895360 / 100 |
| CP2-A2P |  | *Coprinellus* sp. MK307658<br>/ 99.84 | - | - |
| CP2-A3C |  | *Acremonium persicinum*<br>JQ599382 / 99.42 | - | - |
| CP2-A3P |  | *Beauveria* aff. *bassiana*<br>MZ618707 / 100 | - | *Xenoacremonium recifei*<br>KM232105 / 89 |
| CP2-A4C |  | *Acremonium persicinum*<br>JQ599382 / 100 | - | - |
| CP2-A4P |  | *Cyphellophora* aff.<br>*pluriseptata* MH063042.1 /<br>91.7 | - | *Cyphellophora* sp.<br>LR814116 / 80% |
| CP2-A5C |  | *Penicillium* aff.<br>*sumatraense* MH864547 /<br>100 | - | - |
| ND1-A1P | Cloudy Days Act, 1821 | *Penicillium steckii*<br>MZ568311 / 100 | - | *Penicillium steckii*<br>MW196656 / 100 |
| ND2-A1P |  | *Cladosporium* sp.<br>OK242741 / 100 | - | *Cladosporium* aff.<br>*oxysporum* KU216745 /<br>99.7 |

\*If the percent identity is less than 80% and query coverage less than 50%, then the result is indicated as “no match.”

Most of the fungi found belong in the Ascomycota (86%), followed by Basidiomycota (14%). Among the resulting orders, the majority belong in Hypocreales (23%; *Acremonium, Beauveria*, and *Purpureocillium*), Eurotiales (18%; *Aspergillus* and *Penicillium*), and Capnodiales (18%; *Cladosporium*). The Basidiomycota was represented by *Coprinellus, Trametes* and an unidentified species of Psathyrellaceae.

### Cellulolytic activity

The results from cellulolytic activity are shown in Table 3. For the 22 isolates tested, the Enzymatic Index (EI) average was 2.45, with *Cyphellophora* aff. *pluriseptata* CP2-A4P isolated from the Political Constitution being the fungus with the greatest cellulolytic activity (4.0 ± 0.3), followed by *Penicillium steckii* ND1-A1P (3.3 ± 0.3), and *Cladosporium* sp. CP1-A2P (3.3 ± 0.1). In contrast, *Purpureocillium lilacinum* 1539-A1P —from the 1539 Guatemalan Series— was the only isolate without cellulolytic activity. The other fungal isolates showed cellulolytic activity above the levels of the control (*Pleurotus ostreatus*), except for *Trametes* CP1-A3C.

**Table 3.** Enzymatic index (EI) registered by fungal isolates (± indicates standard deviation based on three replicates).

| Isolate code | Taxonomy | Enzymatic Index (EI) |
| --- | --- | --- |
| CP2-A4P | *Cyphellophora* aff. *pluriseptata* | 4.0 ± 0.3 |
| ND1-A1P | *Penicillium steckii* | 3.3 ± 0.3 |
| CP1-A2P | *Cladosporium* sp. | 3.3 ± 0.1 |
| CP2-A1P | *Purpureocillium lilacinum* | 3.2 ± 0.0 |
| 1549-4A1C | Herpotrichiellaceae | 3.1 ± 0.4 |
| CP2-A2C | *Pestalotiopsis kenyana* | 3.1 ± 0.5 |
| CP2-A3P | *Beauveria* aff. *bassiana* | 3.1 ± 0.3 |
| CP1-A1P | *Cladosporium* sp. | 3.0 ± 0.1 |
| CP2-A5C | *Penicillium* aff. *sumatraense* | 2.9 ± 0.7 |
| CP2-A1C | Pleosporales | 2.8 ± 0.2 |
| AI3-A1P | *Aspergillus hiratsukae* | 2.8 ± 0.0 |
| 1549-1A1P | *Penicillium compactum* | 2.6 ± 0.9 |
| AI1-A1C | *Cladosporium* sp. | 2.6 ± 0.1 |
| CP1-A1C | *Periconia* sp. | 2.5 ± 0.3 |
| ND2-A1P | *Cladosporium* aff. *oxysporum* | 2.5 ± 0.2 |
| CP1-A3P | Psathyrellaceae | 2.2 ± 0.2 |
| CP2-A2P | *Coprinellus* sp. | 2.0 ± 0.3 |
| CP2-A4C | *Acremonium persicinum* | 1.8 ± 0.1 |
| CP1-A2C | *Neopestalotiopsis clavispora* | 1.7 ± 0.1 |
| CP2-A3C | *Acremonium persicinum* | 1.6 ± 0.2 |
| Control | *Pleurotus ostreatus* | 1.3 ± 0.1 |
| CP1-A3C | *Trametes hirsuta* | 1.1 ± 0.0 |
| 1539-A1P | *Purpureocillium lilacinum* | 0 |

## Discussion

In this work we determined the chemical and microbiological composition of five important historic documents of Costa Rica, including the Independence Act of 1821. Through spectral techniques, we determined that for the documents dated between 1500 and 1900 (i.e., the Cloudy Days Act, the Independence Act of Costa Rica of 1821, and two documents from the Guatemalan Series of 1539 and 1549), the composition of the paper was cotton (i.e., approximately 90% cellulose (Felgueiras *et al*. 2021), except the second page of the Cloudy Days Act, which is composed of cellulose acetate. The 1991 replicate of the Political Constitution of 1949 was made of cellulose and lignin; this paper presented the greatest amount of lignin, which indicates a modern paper made from wood (hereafter referred to as wood cellulose-based paper).

Despite that modern paper is relatively stable, deterioration is common with increased levels of humidity and acidification produced by oxidation and the presence of microorganisms (Proniewicz *et al*. 2001). Although the 1991 replicate of the Political Constitution from 1949 is the most recent, 15 fungal isolates were obtained, whereas from all the oldest documents (made mainly of cotton), seven in total were obtained. These data suggest that wood-cellulose-based paper possesses characteristics that are more suitable for fungal colonization than cotton-based documents. These observations make sense if we consider the differences between the chemical composition of cotton and paper obtained from other plant fibers such as wood. Cotton contains approximately 90% cellulose (hence it is sometimes referred to as highly pure cellulose), whereas other natural fibers, such as wood, contain 40–55% cellulose combined with other constituents such as lignin and hemicelluloses (Felgueiras *et al*. 2021). Cellulose is considered a two-phase material, having both crystalline and amorphous phases (Ling *et al*. 2019). Cotton cellulose fibers are reported to have a greater degree of polymerization and crystallinity, which generates stronger fibers and greater resistance to hydrolysis and biodegradation. (Itävaara *et al*. 1999). Therefore, these two characteristics (higher cellulose content and higher crystallinity) make cotton a substrate that is less prone to microbial colonization than a material such as wood-based paper, which is the case of the paper of the 1991 replicate of the Political Constitution of 1949. Enzymatically, the reduced ability for microbial colonization of cotton-based papers is related to the more limited access of the cellulase enzyme complex to the substrate due to the orderly and compact architecture of the crystalline cellulose present in cotton (Arantes and Saddler 2010).

In Fig. 1C, the orangish spots over the Independence Act surface indicate oxidation from cellulose and iron, probably caused by both abiotic and biotic factors (Choi 2007). Those areas are fluorescent under UV light and appear as dark spots in UV reflectance photographs (Supplementary Fig. S1). Although excessive humidity can itself trigger oxidation, contributing to the damage of important documents, the inks used can also be affected by this abiotic factor, producing the migration of metal ions of common ink components such as iron and copper, compromising the preservation of historic and cultural heritage due to weakened paper (Henniges *et al*. 2006). Our work determined that the ink employed in the documents from 1821 was ferrogallic, formed by iron sulfate salts in combination with gallic and tannic acids. The last was confirmed from the XRF and multispectral photography (see Supplementary Fig. S1). Ink of this kind is visible under infrared and, in darkness, under UV light (Havermans *et al*. 2003). The implementation of optical spectroscopy techniques, such as those used in this work, have been shown to help identify the early stages of document damage by microorganisms such as fungi, relating changes in the spectral composition to the active presence of fungi (Povolotckaia *et al*. 2019).

In the historic documents, we obtained a total of 22 fungi belonging to 14 genera, of which five (35%) were previously identified in paper-based historic documents (Bensch *et al*. 2018; Pinheiro *et al*. 2019; Romero *et al*. 2021; Trovão and Portugal 2021). The presence of two Chaetothyriales members (isolates 1549-4A1C and CP2-A4P) is interesting because they are inhabitants of environments with limited resources, such as rocks, insects, and ant nests; some species in the order can even become pathogenic for humans, especially in tropical regions ( Attili-Angelis *et al*. 2014; Ahmed *et al*. 2021). *Cyphellophora*, also Chaetothyriales, is closely related to *Phialophora* (Feng *et al*. 2014), which includes species that grow in extremely acidic conditions and have been reported to produce ß-mannanase and ß-glucanase enzymes (Zhao *et al*. 2010; Zhao *et al*. 2012). Our isolate CP2-A4P had the greatest enzymatic index value (4.0 ± 0.3), for which further analysis involving enzyme identification could yield intriguing results. Isolate *Penicillium steckii* ND1-A1P originated from a cotton-based substrate and presented an enzymatic index of 3.3 ± 0.3. This corresponds to what is commonly observed because *Aspergillus* and *Penicillium* species are registered continuously in paper biodeterioration and are known to break the hydrogen bonds, which translates into a weakening of the documents, regardless of their composition (Povolotckaia *et al*. 2019). In total, three isolates corresponding to *Penicillium* were obtained from the Cloudy Days Act, the Political Constitution, and the 1549 Guatemalan Series. We registered only a single isolate corresponding to *Aspergillus hiratsukae* (AI3-A1P), which was recovered from the Independence Act of 1821. This was unexpected because *Aspergillus* spp. are frequent biodeteriogens in cultural heritage objects and spaces in which historic documents are preserved, such as the National Archive of Cuba, in which *Aspergillus*, *Cladosporium*, and *Penicillium* were the most frequent airborne genera reported (Borrego and Perdomo 2016). In this work, *Cladosporium* isolates were present in the Cloudy Days Act (isolate ND2-A1P), the Independence Act (isolate AI1-A1C), and the Political Constitution (isolates CP1-A1P and CP1-A2P), corresponding to cellulose acetate, cotton, and wood cellulose substrates. *Cladosporium* spp. are reported as colonizers in agricultural waste (Herculano *et al*. 2011), artworks (Coronado-Ruiz *et al*. 2018), repositories of historic documents (Borrego and Perdomo 2016), and as extremophiles in high-altitude tropical glaciers (Calvillo-Medina *et al*. 2020).

*Neopestalotiopsis clavispora* (CP1-A2C) and *Pestalotiopsis kenyana* (CP2-A2C) (Xylariales), both obtained from the Political Constitution, belong to a genus known as a common endophyte, saprotroph, and phytopathogen in diverse plants and climates (Reddy *et al*. 2016). Because of the various ecological relationships this genus has with plants, it has attracted attention for its cellulolytic abilities, with over 400 possible enzymes found through the study of a *Pestalotiopsis* isolated from a mangrove (Arfi *et al*. 2013). So far, there are some cellulolytic enzymes known from this genus that have been studied, such as the xylanase (Koh *et al*. 2021) or the cellulases (Goukanapalle *et al*. 2020). Another single isolate obtained from the Political Constitution was *Periconia* CP1-A1C, which belongs to the dark septate fungi Pleosporales, with some isolates reported with thermostable β-Glucosidases suitable for biotechnological processes (Harnpicharnchai *et al*. 2009). *Periconia* species have been encountered in extreme environments, such as deserts, seas, tropical glaciers (Calvillo-Medina *et al*. 2020), and even a new species growing over lithographs from the 19th century (Coronado-Ruiz *et al*. 2018), among others.

Hypocreales isolates were only found in the Political Constitution. Intriguing were two isolates known as insect pathogens, i.e., *Beauveria* (CP2-A3P) and *Purpureocillium* (1539-A1P and CP2-A1P; Shrestha *et al*. 2019). The cellulolytic activity recorded for our *Beauveria* isolate was 3.1 ± 0.3, which is consistent with reports of cellulolytic activity driven by a thermally stable β-Glucosidase in *Beauveria bassiana* (Borgi and Gargouri 2016). Screening for cellulolytic activity in this entomopathogenic species is not commonly done because it is mainly studied for its biological control abilities in multiple agricultural crops (Posada and Vega 2006; Sanjuan *et al*. 2014; Mwamburi 2021). *Beauveria* has also been found as an endophyte, along with *Purpureocillium* (Kepler *et al*. 2013). An isolate of *Purpureocillium lilacinum* has been reported as a biodeteriogen of indoor materials able to grow in alkaline materials, producing damage in limestones and plasters of cultural heritage in Russia (Ponizovskaya *et al*. 2019). The tolerance to extreme conditions by some *Purpureocillium* spp. results in the ability to become pathogenic to humans and resistant to fungicides (Calvillo-Medina *et al*. 2020). Although one isolate recovered from the 1539 Guatemalan Series (cotton-based) was unable to grow in the carboxymethyl cellulose media, isolate CP2-A1P from the Political Constitution (wood cellulose-based) attained EI of 3.2. *Acremonium* isolates CP2-A3C and ND2-A1P registered enzymatic indexes < 3. Previous researchers reported a xylanase produced by *Acremonium cellulolyticus* (Watanabe *et al*. 2014); another group subsequently revised the taxonomy and concluded a misidentification of an *Acremonium* species, reidentifying isolate Y-94 from Japan as *Talaromyces* (Eurotiales) (Fujii *et al*. 2014). Apart from the latter, only a xylanase has been reported for *A. alcalophilum* (Šuchová *et al*. 2020).

Small enzymatic indexes also occurred for three basidiomycetes recovered from the Political Constitution (CP1-A3C, CP1-A3P, and CP2-A2P). These were identified as belonging to Psathyrellaceae, including *Coprinellus* (CP2-A2P). From these, only one report exists of a xylanase produced by *Coprinellus disseminatus*, which was tolerant to varied pH and temperatures (Agnihotri *et al*. 2010). Isolate *Trametes* CP1-A3C registered an EI less than the control *(Pleurotus ostreatus*). Species with tough fruiting bodies, such as *Trametes maxima*, are reported to possess laccase activity, even when exposed to herbicides (Cupul *et al*. 2014); an isolate of *T. versicolor* is reported to cause the effective degradation of fungicides (Rodríguez-Rodríguez *et al*. 2012).

## Conclusions

Regardless of the material of a document of origin —cotton or wood cellulose— most recovered fungal isolates presented cellulolytic activity. From the historic documents sampled, the Political Constitution had the greatest number of isolates, which suggests that wood cellulose-based paper possesses characteristics more suitable for fungal colonization than the oldest cotton-based documents (i.e., documents from 1500–1900). Cotton contains 90% cellulose and has great crystallinity, which makes it more difficult for the enzymatic machinery of microorganisms to degrade these polymers and use them for their nutritional requirements. Even though the oldest documents (e.g., Independence Act and the Cloudy Days Act) yielded few isolates, restoration and improvement of the conditions they are stored in should be implemented to avoid oxidation and weakening of their fibers which could then increase microbiological contamination. The results of our work provide valuable information for establishing the appropriate protocols to undertake that restoration and conservation. Determining the chemical composition of the paper and the composition of the inks, as well as the microbiological load, allows for identification of the most appropriate strategies and treatments to restore documents as important to Costa Rica as the Act of Independence itself. Because historic documents can be considered microhabitats with limited resources, the screening of species with novel biotechnological applications in such environments is a promising and fascinating field. The study of how best to conserve historic documents is vital to preserve, in a satisfactory condition, important sources and records of human history. Multidisciplinary approaches such as the present work can help curators make the best choice of restoration techniques and eventually fulfill the Korean phrase: “Silk can stand five hundred years, but paper can stand one thousand” (Jeong et al., 2014).

## Supporting information

Figure S1. Multispectral photograph of page 127 of the Act of Independence.

Fig. S2. Fungal isolates recovered from the historical documents from the NACR

