## Supplementary information for "Fungi with history: Unveiling the mycobiota of historic documents of Costa Rica"

^1^Centro Nacional de Innovaciones Biotecnológicas (CENIBiot), CeNAT-CONARE, 1174-1200 San José, Costa Rica. 2Centro de Investigaciones en Productos Naturales (CIPRONA), Universidad de Costa Rica, 11501-2060 San José, Costa Rica. ^3^Escuela de Química, Universidad de Costa Rica, 11501-2060 San José, Costa Rica. ^4^Archivo Nacional de Costa Rica, San José, Costa Rica. ^5^Centro de Investigación en Ciencia e Ingeniería de Materiales (CICIMA), Universidad de Costa Rica, 2060 San Pedro, San José, Costa Rica. ^6^Escuela de Biología, Universidad de Costa Rica, 11501-2060 San José, Costa Rica.

*Corresponding authors:

Priscila Chaverri

Escuela de Biología and Centro de Investigaciones en Productos Naturales, Universidad de Costa Rica, 11501-2060 San José, Costa Rica.

Max Chavarría

Escuela de Química and Centro de Investigaciones en Productos Naturales, Universidad de Costa Rica, 11501-2060 San José, Costa Rica.

**LEGENDS OF SUPPLEMENTARY FIGURES**

**Figure S1. Multispectral photograph of page 127 of the Act of Independence.** **A**. Reflectance ultraviolet photograph **B**. Fluorescence ultraviolet photograph. **C**. Visible photograph. Surface indicate an oxidation process from the cellulose and iron, probably caused by both abiotic and biotic factors.

**Figure S2. Fungal isolates recovered from the historical documents from the NACR**. **A.** 1539-A1P, *Purpureocillium lilacinum* **B.** 1549-1A1P, *Penicillium compactum*. **C.** 1549-4A1C, Herpotrichiellaceae. **D.** AI1-A1C, *Cladosporium* sp. **E.** AI3-A1P, *Aspergillus hiratsukae* **F.** CP1-A1C, *Periconia* sp. **G.** CP1-A1P, *Cladosporium* sp. **H.** CP1-A2C, *Pestalotiopsis microspora* **I.** CP1-A2P, *Cladosporium* sp. **J.** CP1-A3C, *Trametes hirsuta* **K.** CP1-A3P, *Unidentified Psathyrellaceae*. **L.** CP2-A1C, Unidentified Pleosporales. **M.** CP2-A1P, *Purpureocillium lilacinum* **N.** CP2-A2C, *Pestalotiopsis trachycarpicola* **O.** CP2-A2P, *Coprinellus* sp. **P.** CP2-A3C, *Acremonium persicinum* **Q.** CP2-A3P, *Beauveria aff. bassiana* **R.** CP2-A4C, *Acremonium persicinum* **S.** CP2-A4P, *Cyphellophora aff. pluriseptata*. **T.** CP2-A5C, *Penicillium aff. sumatraense* **U.** ND1-A1P, *Penicillium steckii* **V.** ND2-A1P, *Cladosporium* sp.
