## Supplementary figures and images for "Fungi with history: Unveiling the mycobiota of historic documents of Costa Rica"

### Fig. S2. Fungal isolates recovered from the historical documents from the NACR

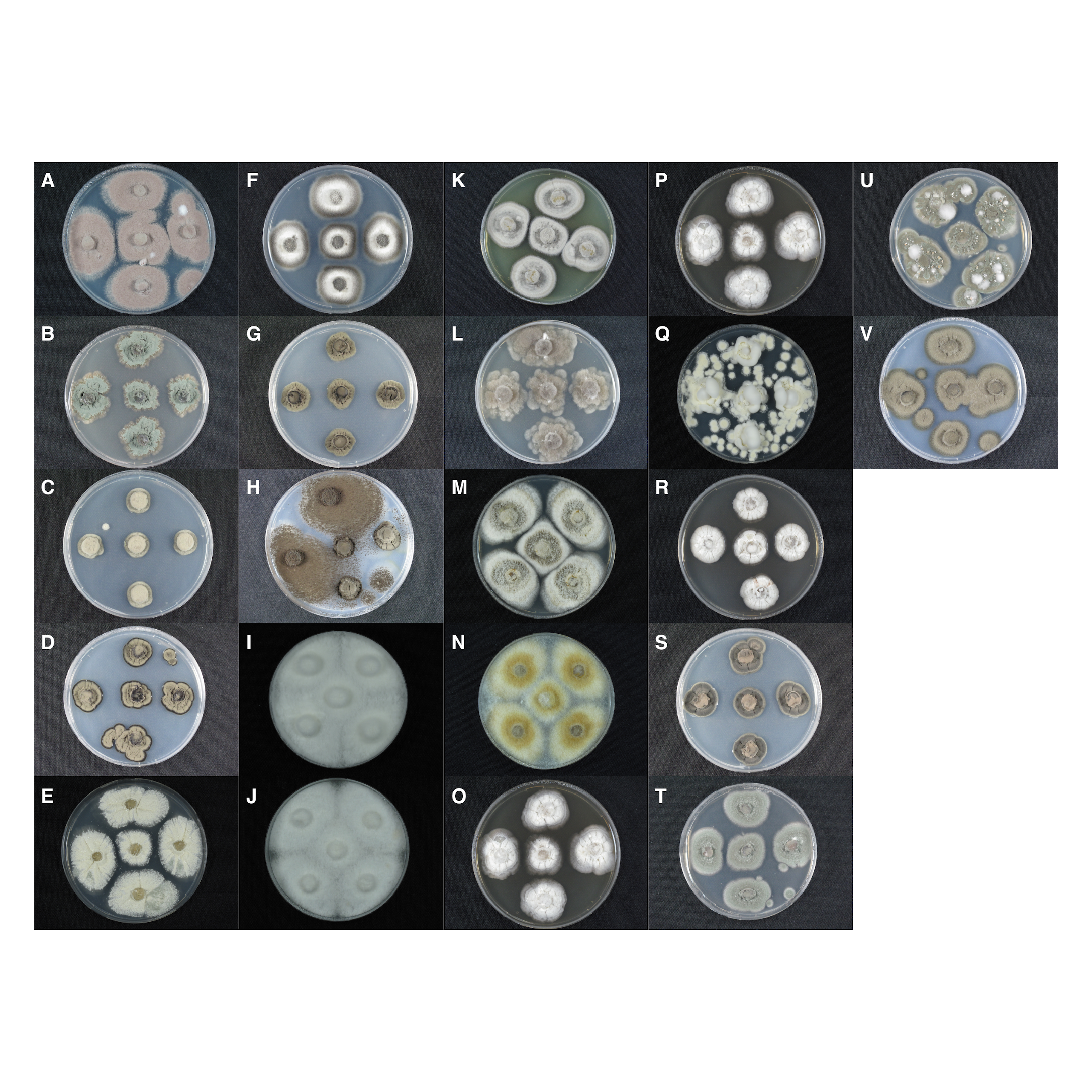

### Figure S1. Multispectral photograph of page 127 of the Act of Independence.

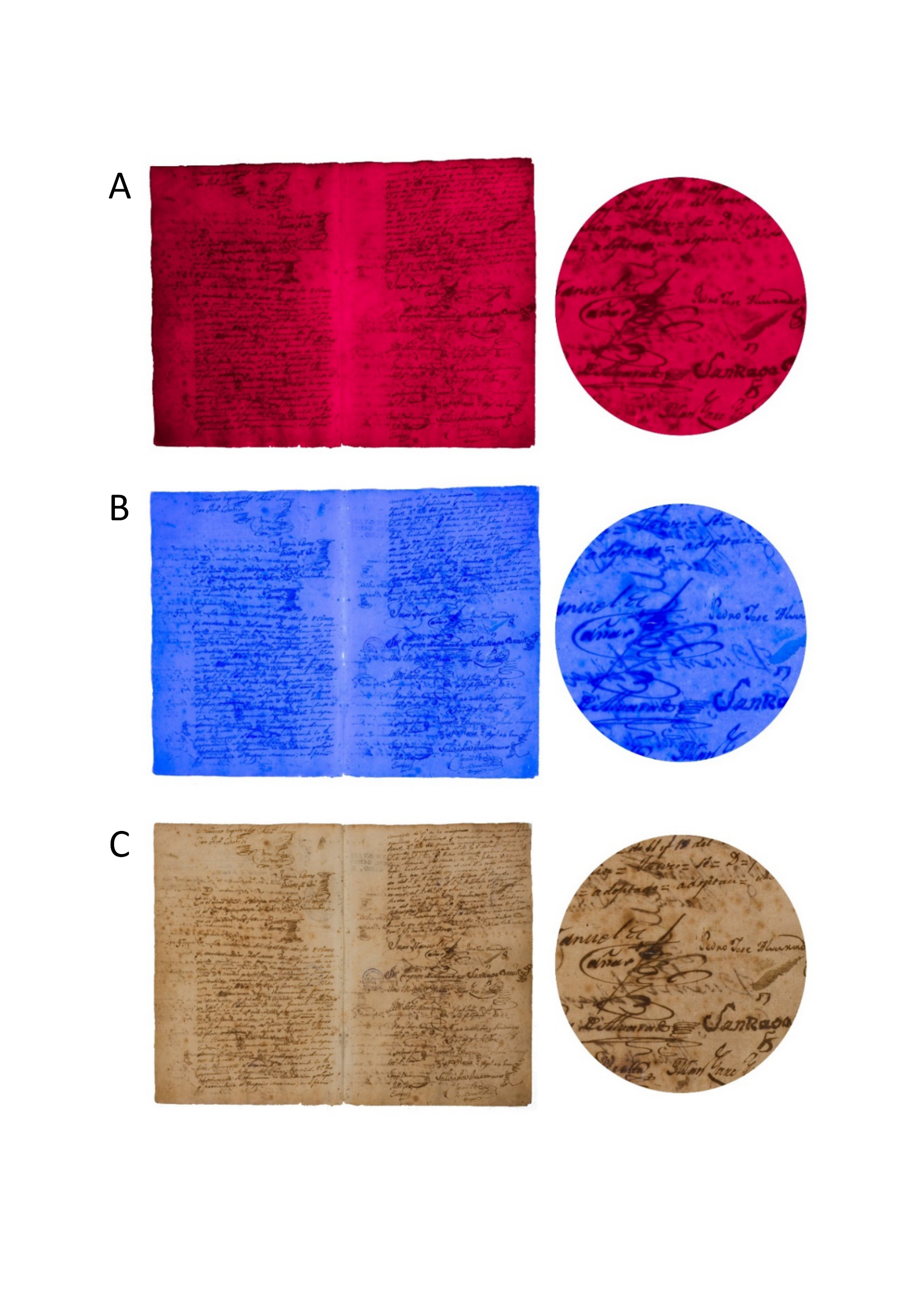
